# Comparison of Shiga toxin-encoding bacteriophages in highly pathogenic strains of Shiga toxin-producing *Escherichia coli* O157:H7 in the UK

**DOI:** 10.1101/843987

**Authors:** Daniel A Yara, David R Greig, David L Gally, Timothy J Dallman, Claire Jenkins

## Abstract

2.

Over the last 35 years in the UK the burden of Shiga toxin-producing *Escherichia coli* (STEC) O157:H7 infection has, during different periods of time, been associated with five different sub-lineages (1983-1995: Ia, I/IIa and I/IIb, 1996-2014: Ic and 2015-2018: IIb). The acquisition of a *stx2a*-encoding bacteriophage by these five sub-lineages appears to have coincided with their respective emergences. Oxford Nanopore Technology (ONT) was used to sequence, characterise and compare the *stx*-encoding prophage harboured by each sub-lineage to investigate the integration of this key virulence factor. The *stx2a*-encoding prophage from each of the lineages causing clinical disease in the UK were all different, including the two UK sub-lineages (Ia and I/IIa) circulating concurrently and causing severe disease in the early 1980s. Comparisons between the *stx2a*-encoding prophage in sub-lineages I/IIb and IIb revealed similarity to the prophage commonly found to encode *stx2c*, and the same site of bacteriophage integration (*sbcB*) as *stx2c* encoding prophage. These data suggest independent acquisition of previously unobserved *stx2a*-encoding phage is more likely to have contributed to the emergence of STEC O157:H7 sub-lineages in the UK than intra-UK lineage to lineage phage transmission. In contrast, the *stx2c*-encoding prophage showed a high level of similarity across lineage and time, consistent with the model of *stx2c* being present in the common ancestor to extant STEC O157:H7 and maintained by vertical inheritance in the majority of the population. Studying the nature of the *stx*-encoding bacteriophage contributes to our understanding of the emergence of highly pathogenic strains of STEC O157:H7.

**Impact statement:** The application of ONT technology to sequence UK epidemic strains of STEC O157:H7 revealed *stx2a*-encoding prophage exhibit a high level of diversity. There was little evidence of geographical or temporal patterns of relatedness, or of intra-UK transmission of *stx2a*-encoding prophage between indigenous strains. The *stx2a*-encoding prophage in the UK lineages associated with severe disease appear to be acquired independently and most likely from different geographical and/or environmental sources. These data provide supporting evidence for the existence of a dynamic environmental reservoir of *stx2a*-encoding prophage that pose a threat public health due to their potential for integration into competent, indigenous sub-lineages of STEC O157:H7. We also provide further evidence that *stx2c*-encoding prophage exhibit a high level of similarity across lineage, geographical region and time, and have likely been maintained and inherited vertically.

**Data summary:** All FASTQ files and assemblies of samples sequenced in this project were submitted to the National Centre for Biotechnology Information (NCBI). All data can be found under BioProject: PRJNA315192 - https://www.ncbi.nlm.nih.gov/bioproject/?term=PRJNA315192. Strain specific details can be found in the methods section under data deposition.

Publicly available data used in this project can be found via Table 1 and data bibliography.

## 5. Introduction

Shiga toxin-producing *Escherichia coli* (STEC) serotype O157:H7 is a zoonotic pathogen that causes gastrointestinal symptoms in humans. A sub-set of patients (mainly children and the elderly) are at risk of developing haemolytic uremic syndrome (HUS), a potentially fatal systemic condition primarily associated with acute renal failure, cardiac and neurological complications [1]. STEC O157:H7 emerged as a public health concern during the early 1980s and was first isolated in the United Kingdom in July 1983 from three cases linked to an outbreak of HUS [2]. Throughout the 1980s, the increasing number of outbreaks of gastrointestinal disease and HUS associated with this serotype, stimulated the development of sub-typing methods that provided a higher level of strain discrimination than serotyping. In the late 1980s, a phage typing scheme, developed by the Canadian Public Health Laboratory Service was adopted by Public Health England (PHE, then the Public Health Laboratory Service) [3], and is still used today. In 2015, PHE implemented whole genome sequencing (WGS) for routine surveillance of STEC O157:H7 in England [4].

The primary STEC virulence factor is the Shiga toxin (Stx), which targets cells expressing the glycolipid globotriaosylceramide, disrupting host protein synthesis and causing apoptotic cell death. Strains of STEC O157:H7 in the UK produce *stx1a, stx2a* and *stx2c*, either individually or in any combination [5]. Strains harbouring *stx2a*, either alone or in combination with *stx1a* and/or *stx2c* are significantly associated with causing severe disease, including HUS [5,6] and are associated with more efficient transmission within the ruminant reservoir [7]. The genes encoding the *stx* subtypes are located on active bacteriophage that can be acquired and integrated into the chromosome of STEC O157:H7 strains. There is evidence that the different prophage backgrounds that harbour *stx* genes can contribute to differential toxin production and may ultimately affect clinical outcome [8].

There are three main lineages of STEC O157:H7 (I, II and I/II) and eight sub-lineages (Ia, Ib, Ic, IIa, IIb, IIc, I/IIa and I/IIb). In the UK, the outbreaks of STEC O157:H7 in the 1980s were caused by strains belonging to sub-lineage Ia (mainly comprising phage type (PT)1 and PT4), sub-lineage I/IIa (comprising PT2), and sub-lineage I/IIb (comprising PT49) [9]. Throughout the 1990s, these three lineages declined and all but disappeared. Concurrently, we observed a dramatic rise of sub-lineage Ic (mainly comprising PT21/28), in addition to a steady increase in the number of cases of sub-lineage IIc (mainly comprising PT8) [5,9]. Since 2012, the number of cases of PT21/28 has declined and an unusual PT8 variant belonging to sub-lineage IIb has emerged [10].

With the exception of sub-lineage IIc (PT8), which is not associated with HUS cases in the UK [5], all the dominant UK sub-lineages over time encode *stx2a*, and the acquisition of a *stx2a*-encoding bacteriophage appears to have coincided with their respective emergences [5,10]. The aim of this investigation was to use Oxford Nanopore Technology (ONT) to sequence, characterise and compare the *stx*-encoding prophage harboured by each of the UK sub-lineages to determine the similarity of the *stx*-encoding prophage acquired by each lineage. Studying the nature of the *stx*-encoding bacteriophage will contribute to our understanding of the emergence of highly pathogenic strains of STEC O157:H7.

## 6. Methods

### Bacterial strains

Six strains of STEC O157:H7 were selected for sequencing from the PHE archive on the basis of being the earliest representative of each of the sub-lineages that acquired the *stx2a*-encoding prophage (Table 1). Eleven publicly available sequences were also included in the analysis for context. Of these, seven originated from the UK, five were the cause of four published outbreaks [11,12,13,14], three were from the USA [15,16] and one from Japan [17] (Table 1).

**Table 1 –.**
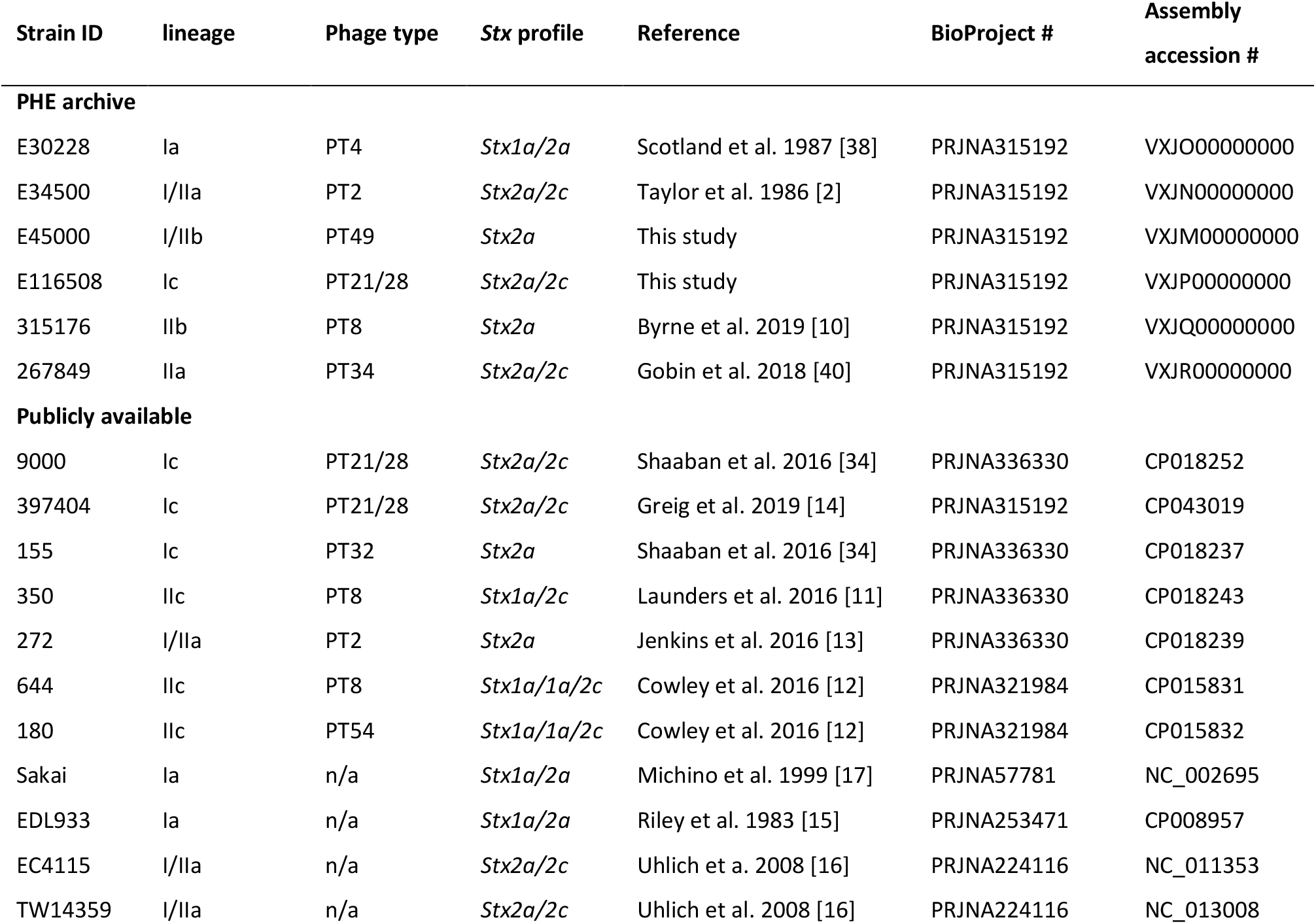
Summary of the PHE archived and publicly available strains used within this study, their Strain ID, lineage, phage type, Stx profile, assembly accession numbers and NCBI BioProject.

### Short read sequencing on the Illumina HiSeq 2500

Genomic DNA (gDNA) was extracted from cultures of STEC O157:H7 using the Qiagen Qiasymphony (Qiagen, Hilden, Germany). The sequencing library was prepared using the Nextera XP kit (Illumina, San Diego, USA) for sequencing on the Illumina HiSeq 2500 (Illumina, San Diego, USA) instrument run with the fast protocol. High quality trimmed (leading and trailing trimming at <Q30 using Timmomatic v0.27 [18].

### Long read sequencing using ONT and data processing

Genomic DNA was extracted and purified using the Qiagen Genomic Tip, midi 100/G (Qiagen, Hilden, Germany) with minor alterations including no vigorous mixing steps (performed by inversion) and elution into 100μl double processed nuclease free water (Sigma-Aldrich, St. Louis, USA). gDNA for each extract was quantified using a Qubit and the HS (high sensitivity) dsDNA Assay Kit (Thermofisher Scientific, Waltham, USA) to manufacturer’s instructions. Library preparation was performed on several instances using both rapid barcoding kits - SQK-RBK00(1/4) and native barcoding kit (SQK-LSK108 and EXP-NBD103) (Oxford Nanopore Technologies, Oxford, UK). The prepared libraries were loaded onto a FLO-MIN106 R9.4.1 flow cells (Oxford Nanopore Technologies, Oxford, UK) and sequenced using the MinION (Oxford Nanopore Technologies, Oxford, UK) for 48 hours.

Data produced in a raw FAST5 format was basecalled and de-multiplexed using Albacore V2.3.3 (Oxford Nanopore Technologies) into FASTQ format and grouped in each samples’ respective barcode. De-multiplexing was performed using Deepbinner v0.2.0 [19]. Run metrics were generated using Nanoplot v1.8.1 [20]. The barcode and y-adapter from each sample’s reads were trimmed, and chimeric reads split using Porechop v0.2.4 [21]. Finally, the trimmed reads were filtered using Filtlong v0.1.1 [22] with the following parameters; min length=1000, keep percent=90 and target bases=550 Mbp, to generate approximately 100x coverage of the STEC genome with the longest and highest quality reads.

### De novo assembly, polishing, reorientation and annotation

Trimmed ONT FASTQ files were assembled using Canu v1.7 [23] and the filtered ONT FASTQ files were assembled using both Unicycler v0.4.2 [24] with the following parameters min_fasta_length=1000, mode=normal and Flye v2.4.2 [25] using default parameters. The assembly for each sample that had the highest N50 and lowest number of contigs with the assembly size (between 5.3-6.0 Mbp) were taken forward. Polishing of the assemblies was performed in a three-step process firstly, using Nanopolish v0.11.1 [26] using both the trimmed ONT FASTQs and FAST5s for each respective sample accounting for methylation using the --methylation-aware=dcm and --min-candidate-frequency=0.5. Secondly, Pilon v1.22 [27] using Illumina FASTQ reads as the query dataset with the use of BWA v0.7.17 [28] and Samtools v1.7 [29]. Finally, Racon v1.2.1 [30] also using BWA v0.7.17 [28] and Samtools v1.7 [29] was used with the Illumina reads for two cycles to produce a final assembly for each of the samples. As the chromosome from each assembly was circularised and closed, they were re-orientated to start at the *dnaA* gene (NC_000913) from *E. coli* K12, using the --fixstart parameter in circlator v1.5.5 [31]. Prokka v1.13 [32] with the use of a personalised database (an amino acid FASTA which included all genes annotated in the publicly available samples used in this study) was used to annotate the final assemblies.

### Prophage detection, excision and processing

Prophages across all samples were detected and extracted using the updated Phage Search Tool (PHASTER) [33]. Prophage extraction from the genome occurred regardless of prophage size or PHASETER quality score and any detected prophages separated by less than 4kbp were conjoined into a single phage using Propi v0.9.0 as described in Shaaban, et al. 2016 [34]. From here the prophages were trimmed to remove any non-prophage genes and were again annotated using Prokka v 1.13 [32] with the use of a personalised database (an amino acid FASTA which included all genes annotated in the publicly available samples used in this study).

### Mash and phylogeny

Mash v2.2 [35] was used to sketch (sketch length 1000, kmer length, 21) the extracted prophages in the samples sequenced in this study and all Shiga toxin encoding prophages found in the publicly available STEC genomes in table 1. The pairwise Jaccard distance between the prophages was calculated and a neighbour joining tree computed and visualised using FigTree v1.4.4 [36].

### Visualisation tools

All gene diagrams were constructed using Easyfig v2.2.3 [37]. Phylogenetic trees were visualized and annotated using FigTree v1.4.4 [36].

### Data deposition

Illumina FASTQ files are available from BioProject: PRJNA315192 under the following SRA accessions E30228: SRR10290290, E34500: SRR10290289, E45000: SRR10290288, E116508: SRS941727, 315176: SRR6051955 and 267849: SRR3742262. Nanopore FASTQ files are available from BioProject: PRJNA315192 under the following SRA accessions, E30228: SRR10103064, E34500: SRR10103063, E45000: SRR10103062, E116508: SRR10103065, 315176: SRR10103066 and 267849: SRR10103067.

Assemblies can be found under BioProject: PRJNA315192 under the following accessions, E30228: VXJO00000000, E34500: VXJN00000000, E45000: VXJM00000000, E116508: VXJP00000000, 315176: VXJQ00000000 and 267849: VXJR00000000.

## 7. Results and Discussion

### Genomic features of the samples sequenced in this study

All six isolates, selected for sequencing from the PHE archive on the basis of being the earliest representative of each of the sub-lineages that acquired the *stx2a*-encoding prophage, assembled into closed chromosomes with one or more plasmids. The isolates belonging to sub-lineage Ia PT4 (E30228) and sub-lineage IIb PT8 (315176) each assembled into a chromosome (5,416,109 bp and 5,579,120 bp, respectively) and two plasmids (Table 2). The sequence data from the other four isolates each assembled into a chromosome of between 5,359,964 - 5,571,891 bp and a single plasmid (Table 2). The pO157 (IncFIB) plasmid was found in all samples sequenced in this study. The number of prophage in each of the genomes of the six isolates varied from 14 in the isolate belonging to sub-lineage I/IIa PT2 to 17 from the isolates belonging to sub-lineages I/IIb PT49 and Ic PT21/28 (Figure 1).

**Figure 1 –.**
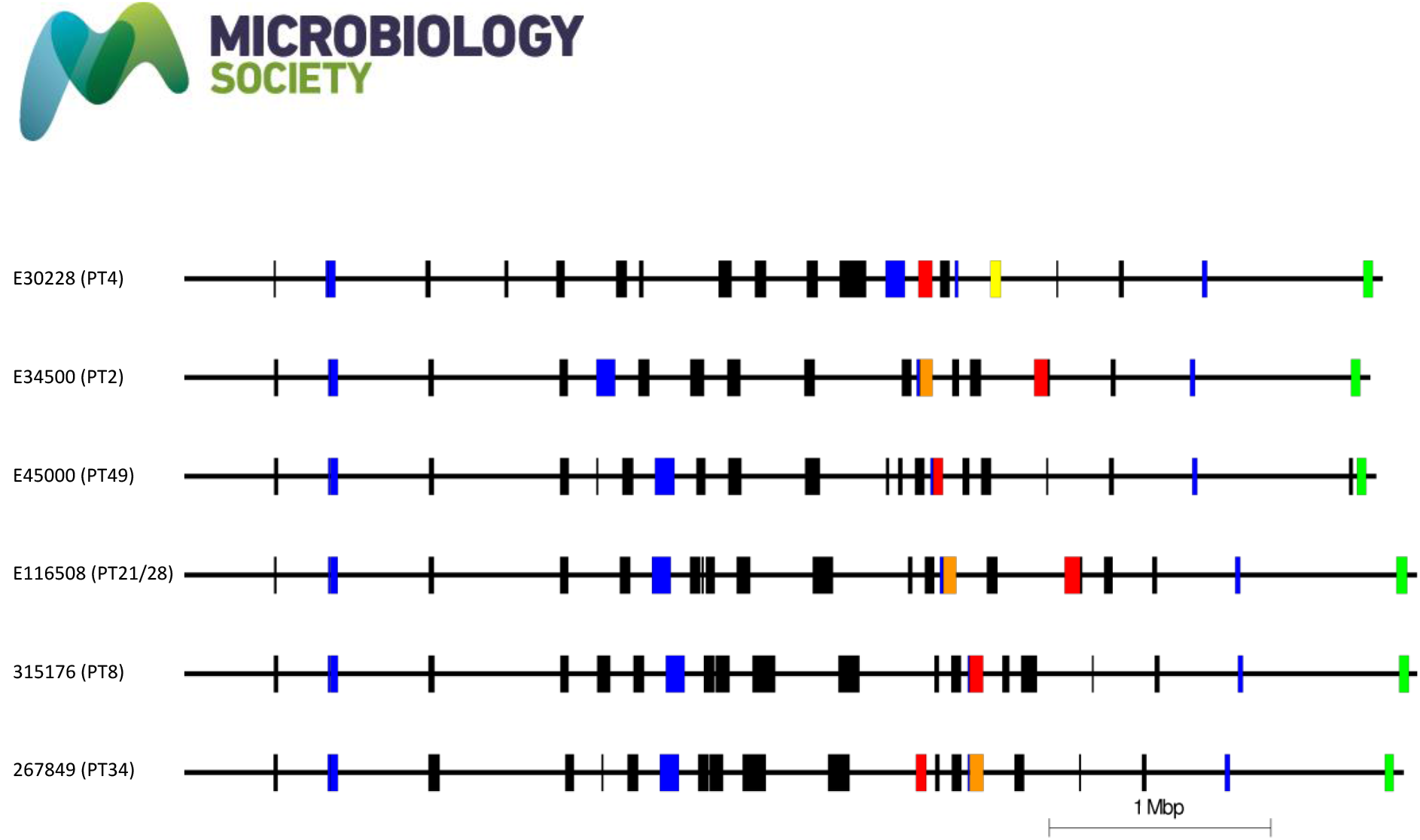
Easyfig diagram representing the chromosome and prophage content within the samples sequenced in this study (Order descending PT4, PT2, PT49, PTPT21/28, PT8 and PT34). *Stx2a*-encoding, *Stx2c*-encoding and *Stx1*-encoding prophages are highlighted in red, orange and yellow respectively. Prophage like elements are coloured blue and the locus of enterocyte effacement in green.

**Table 2 –.**
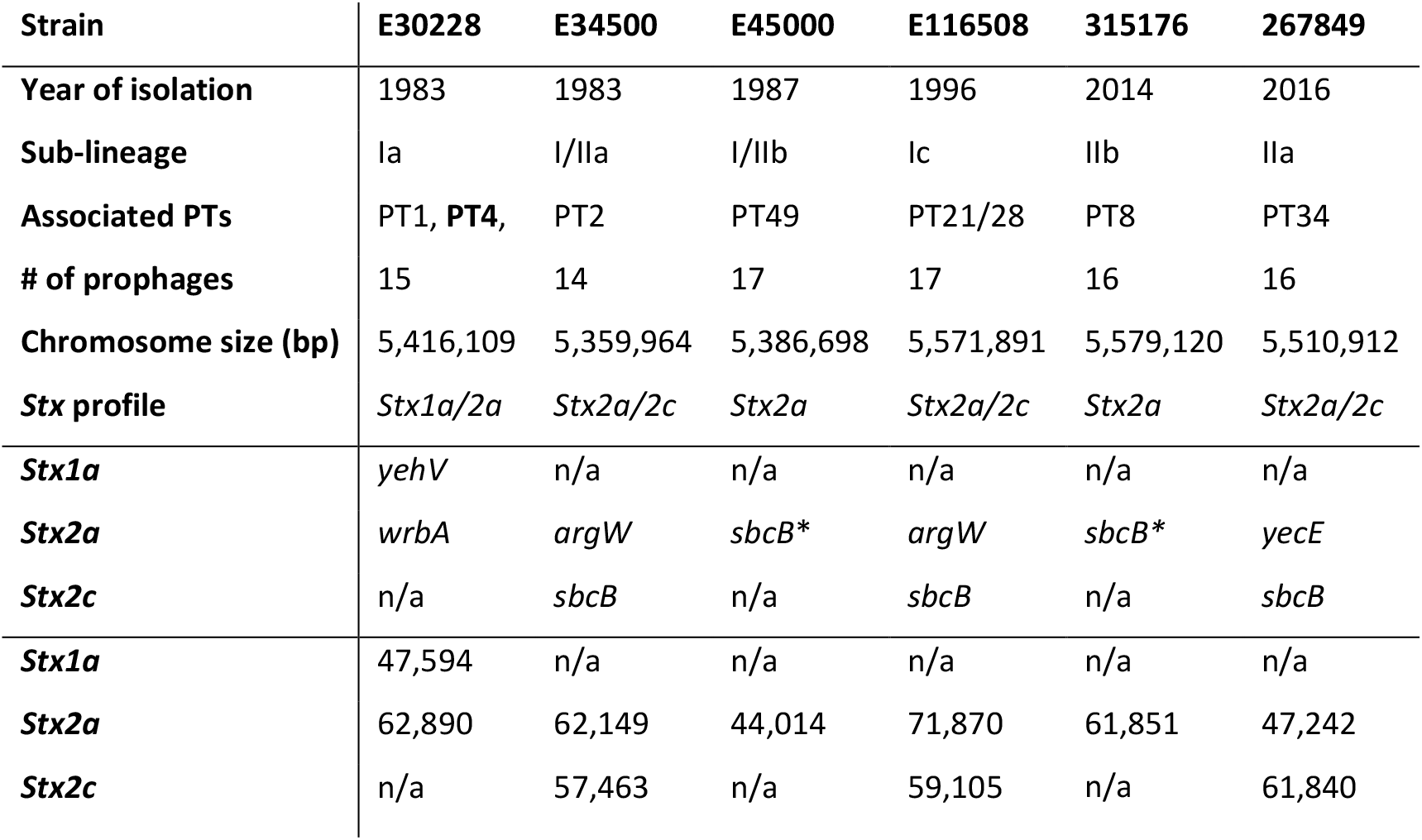
Table showing all strains sequenced in this study, their year of isolation, sub-lineage, phage type, *Stx* profile, the site of bacteriophage insertion (SBI) and prophage size. * denotes *stx2a* genes in a *stx2c* associated prophage structure.

| Strain | E30228 | E34500 | E45000 | E116508 | 315176 | 267849 |
| --- | --- | --- | --- | --- | --- | --- |
| Year of isolation | 1983 | 1983 | 1987 | 1996 | 2014 | 2016 |
| Sub-lineage | Ia | I/IIa | I/IIb | Ic | IIb | IIa |
| Associated PTs | PT1, PT4, | PT2 | PT49 | PT21/28 | PT8 | PT34 |
| # of prophages | 15 | 14 | 17 | 17 | 16 | 16 |
| Chromosome size (bp) | 5,416,109 | 5,359,964 | 5,386,698 | 5,571,891 | 5,579,120 | 5,510,912 |
| Stx profile | *Stx1a/2a* | *Stx2a/2c* | *Stx2a* | *Stx2a/2c* | *Stx2a* | *Stx2a/2c* |
| *Stx1a* | *yehV* | n/a | n/a | n/a | n/a | n/a |
| *Stx2a* | *wrbA* | *argW* | *sbcB\** | *argW* | *sbcB\** | *yecE* |
| *Stx2c* | n/a | *sbcB* | n/a | *sbcB* | n/a | *sbcB* |
| *Stx1a* | 47,594 | n/a | n/a | n/a | n/a | n/a |
| *Stx2a* | 62,890 | 62,149 | 44,014 | 71,870 | 61,851 | 47,242 |
| *Stx2c* | n/a | 57,463 | n/a | 59,105 | n/a | 61,840 |

### Comparison of the stx1a-encoding prophage

Six of the isolates analysed in this study, contained a prophage encoding *stx1a* (Table 1 and Figure 3). The *stx1a*-encoding prophage from the isolate belonging to sub-lineage Ia PT4 (E30228), among the first to be isolated in the UK in 1983, shared similarity with *stx1a*-encoding prophage found in EDL933 and Sakai, two international outbreak strains that also belonged to sub-lineage Ia (Table 1 and Figures 2 and 3). EDL933 caused an outbreak in USA in 1982 linked to contaminated hamburgers [15] and was temporally but not geographically linked to the UK isolate. The outbreak in Sakai City, Japan, associated with contaminated radish sprouts occurred in 1996 [17], and was both temporally and geographically distinct from EDL933 and E30228 (Figures 2 and 3). Previous analysis of isolates of sub-lineage Ia harbouring *stx1a*-encoding prophage indicate the *stx1a* prophage is likely ancestral and inherited vertically [5]. This is consistent with the strains analysed in this study encoding a similar *stx1a* prophage, despite being isolated at different times and geographical locations.

**Figure 2 –.**
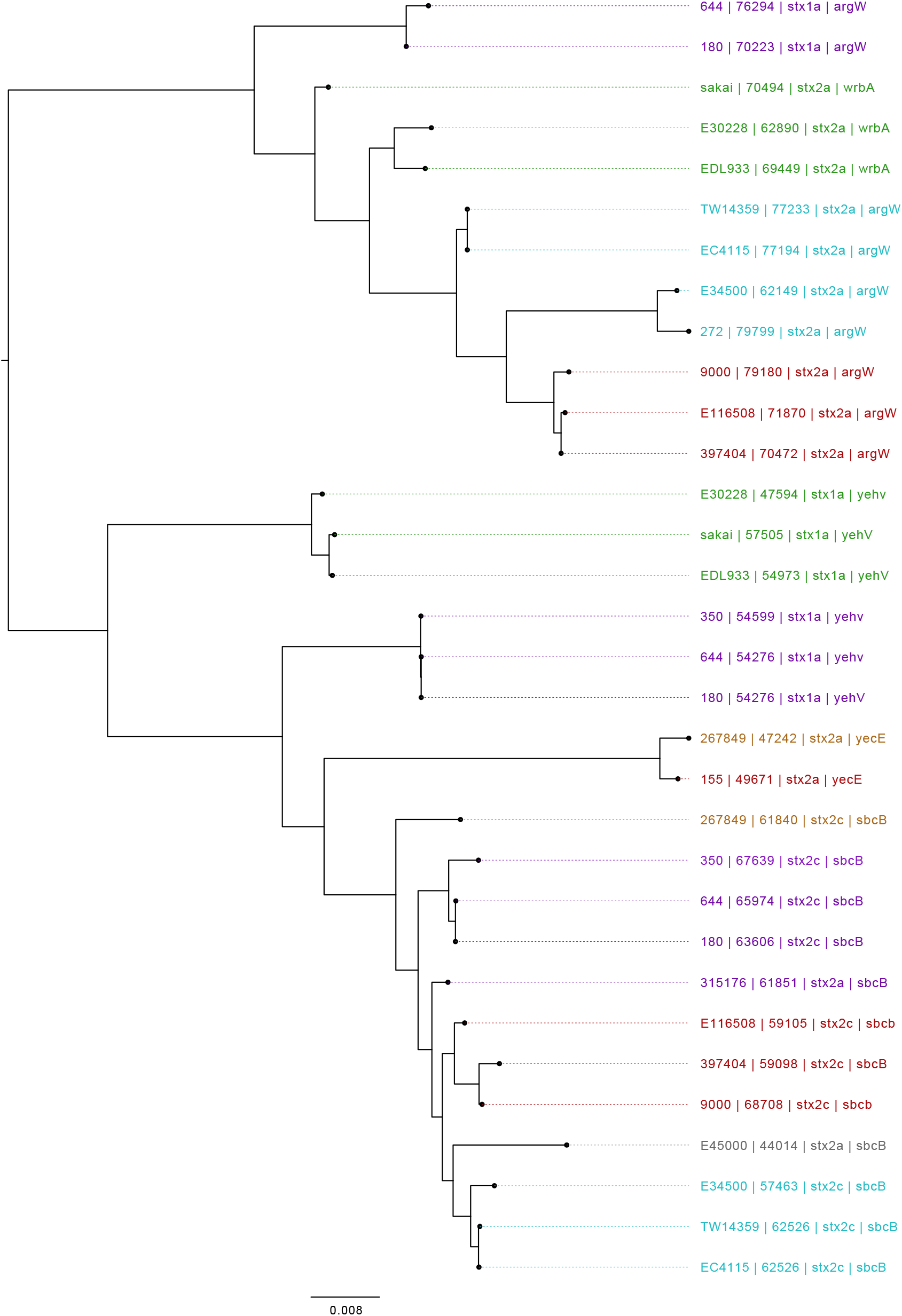
Mid-rooted tree of Shiga toxin-encoding prophages based on Jaccard distance produced from Mash. Strains are annotated as Strain ID, length, *Stx* profile and SBI. Strains are coloured by sub-lineage – Green: Ia, Red: Ic, Blue: I/IIa, Grey: I/IIb, Orange: IIa, Pink: IIb Purple: IIc.

**Figure 3 –.**
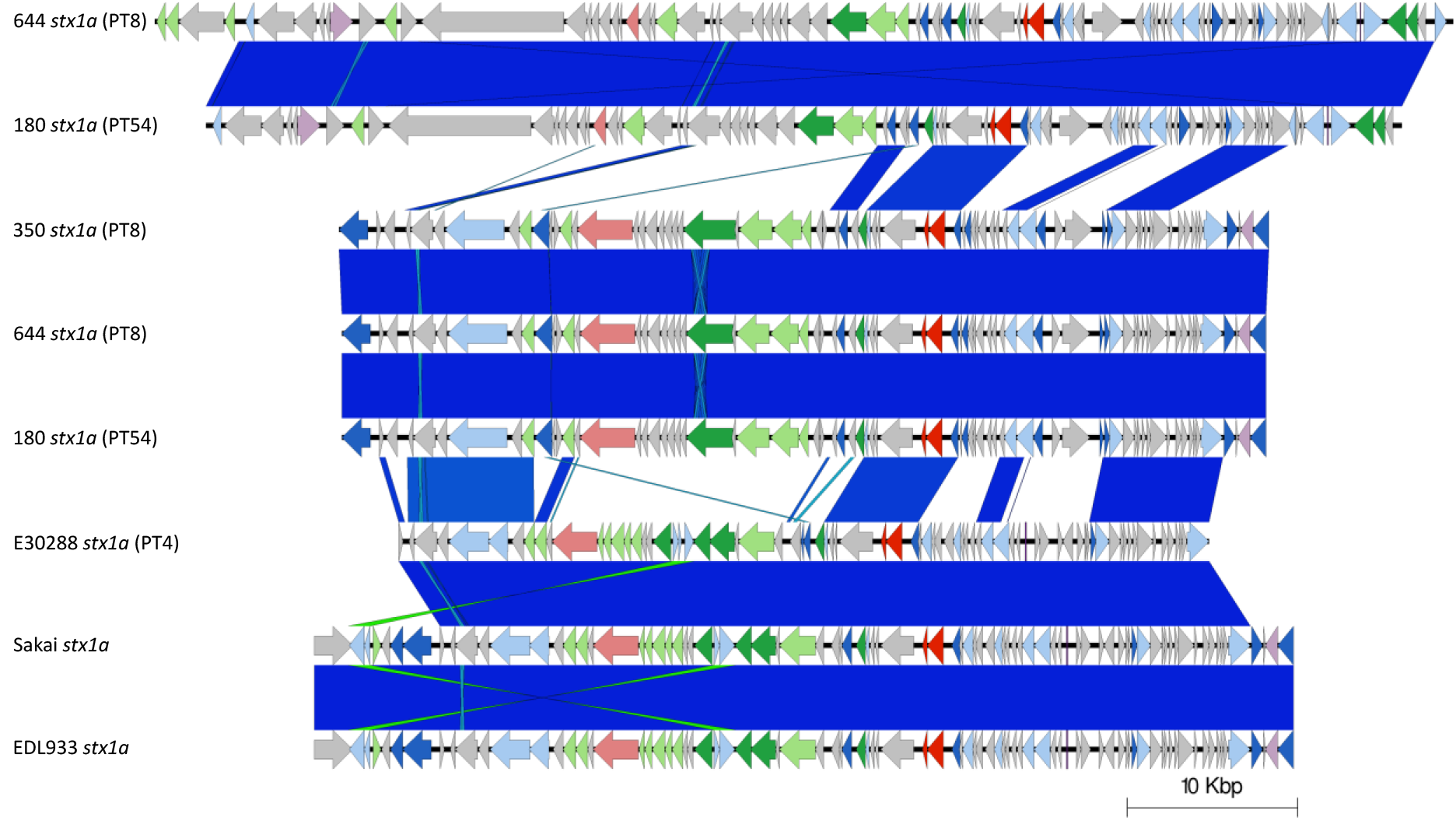
Easyfig plot comparing the *stx1a*-encoding prophages from 644 (x2), 180 (x2), 350, E30288, EDL933 and Sakai. Arrow indicates gene direction. Recombination/replication genes shown in light blue, regulation associated genes in dark blue. Effector genes shown in pink, structure and lysis associated genes shown in light and dark green respectively and tRNAs shown as purple lines, finally grey are hypothetical genes.

The *stx1a*-encoding prophages from three isolates belonging to sub-lineage IIc associated with foodborne outbreaks in the UK [11,12] cluster together based on Mash distance but were distinct from the *stx1a*-encoding prophages harboured by the sub-lineage Ia strains described above. As previously described by Shaaban et al. (2017) [34], two of these strains (664 PT8 and 180 PT54) linked to a foodborne outbreak in Northern Ireland in 2013 [12] had an additional but different *stx1a*-encoding prophage within the same chromosome (Figure 3). Therefore, three different *stx1a*-encoding prophages in two different lineages (Ia and IIc), were identified in this study (Figures 3 and 4).

**Figure 4 –.**
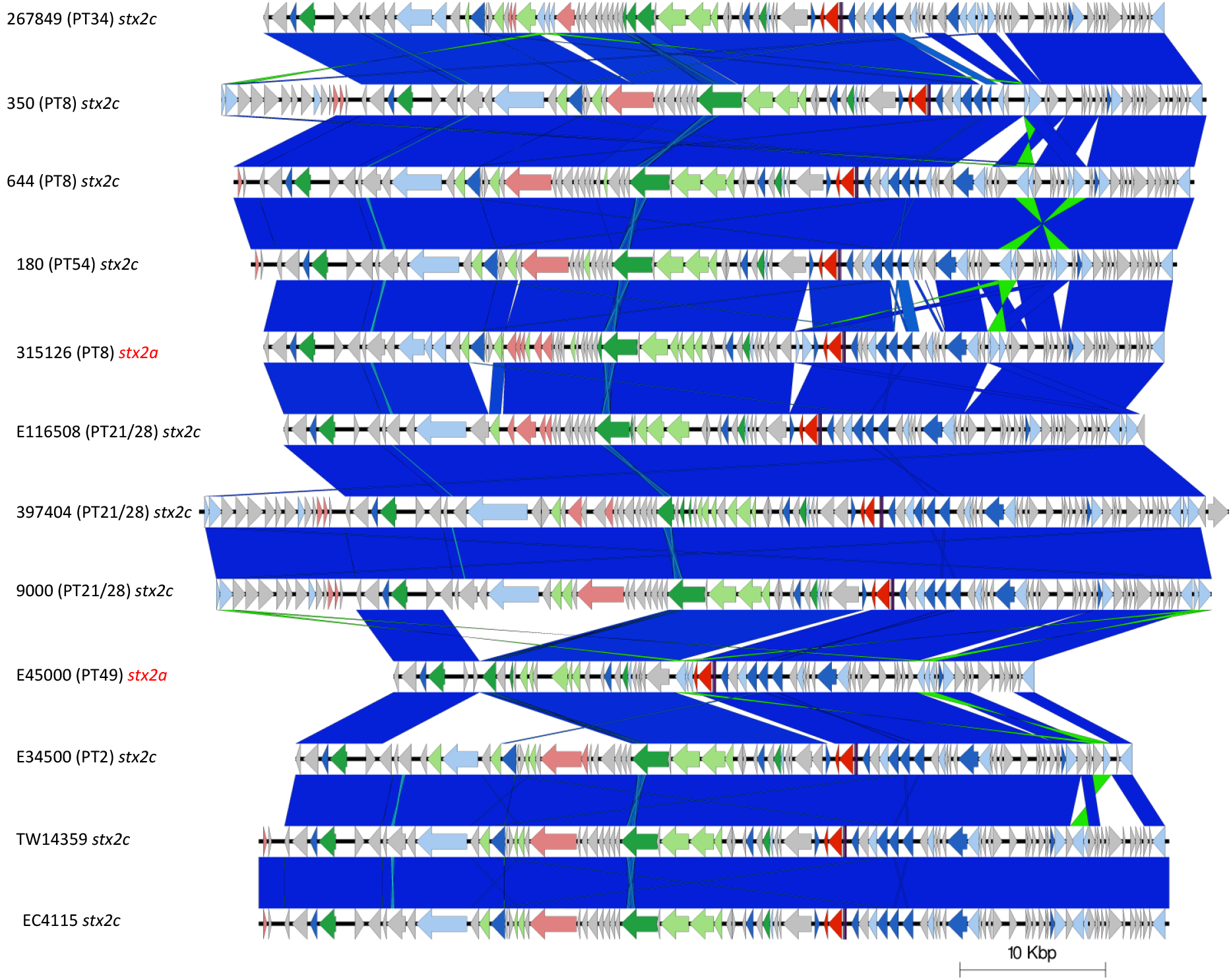
Easyfig plot comparing the *stx2c*-encoding prophages from all samples in the study, including two *stx2a* prophages are in a *stx2c* associated prophage structure (315126 and E45000). Arrow indicates gene direction. Recombination/replication genes shown in light blue, regulation associated genes in dark blue. Effector genes shown in pink, structure and lysis associated genes shown in light and dark green respectively and tRNAs shown as purple lines, finally grey are hypothetical genes.

### Comparison of stx2c-encoding prophage

Nine isolates from four different sub-lineages (Ic, I/IIa, IIa and IIc) contained *stx2c*-encoding prophage. The *stx2c*-encoding prophage from each isolate clustered together based on Mash distance and also aligned across the length of the prophage with few structural variations (Figure 4). The *stx2c* prophage from strains within the same sub-lineage were more similar based on Mash distance then *stx2c* prophage in strain from different lineages (Table 1 and Figures 3 and 5). These strains were isolated over a wide time frame from 1983 to 2016 and from strains isolated in different countries including the UK, Ireland and the USA providing further evidence that *stx2c*-encoding prophage show a high level of similarity across lineage, time and geographical region [34] (Table 1). This is consistent with the model of *stx2c* being present in the common ancestor to extant STEC O157:H7 and maintained by vertical inheritance in the majority of the population.

**Figure 5 –.**
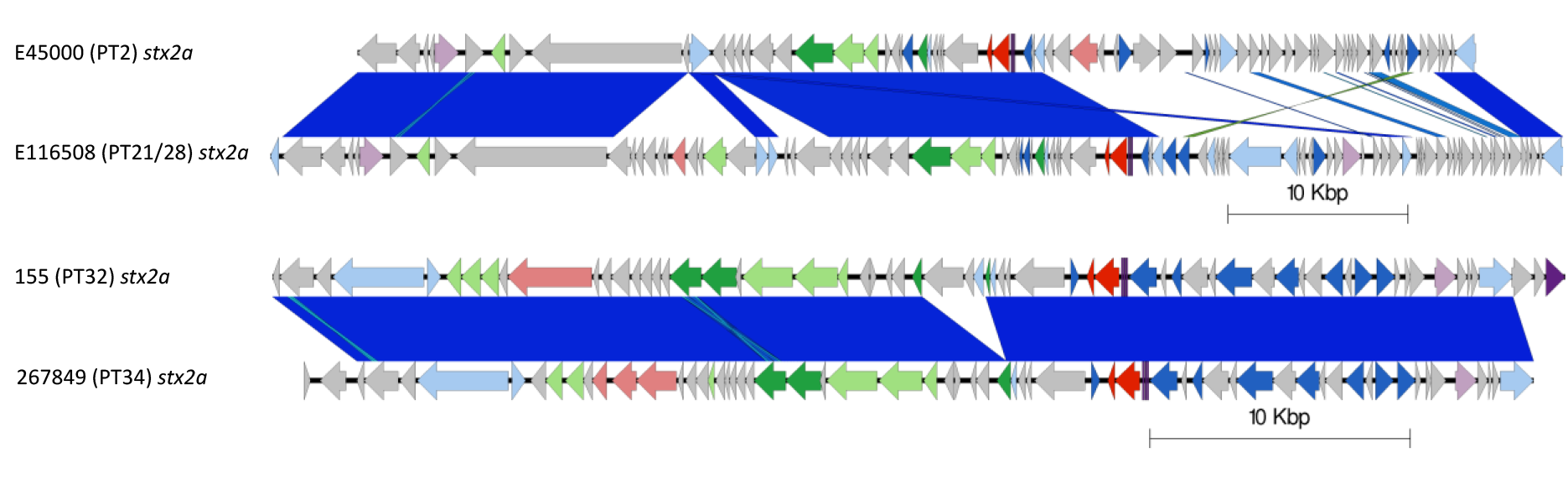
Two Easyfig plots comparing the *stx2a*-encoding prophages from E45000 with E116508 (above) and 155 and 267849 (below) order descending. Arrow indicates gene direction. Recombination/replication genes shown in light blue, regulation associated genes in dark blue. Effector genes shown in pink, structure and lysis associated genes shown in light and dark green respectively and tRNAs shown as purple lines, finally grey are hypothetical genes.

### Comparison of stx2a-encoding prophage

Certain strains that shared lineage, PT and geography harboured similar *stx2a*-encoding prophage. Examples included (i) the two sub-lineage Ic PT21/28 isolates from the UK, (ii) the two sub-lineage I/IIa PT2 isolates from the UK and (iii) the two isolates from sub-lineage I/lla from the USA (Table 1 and Figures 2 and 5). Isolates designated E30228 and EDL933, both sub-lineage Ia and temporally related but geographically distinct, also encoded similar *stx2a*-encoding prophage (Table 1 and Figures 2 and 5), as did isolates 155 (sub-lineage Ic PT32) and 267849 (sub-lineage IIa PT34), which were unrelated temporally and geographically.

Compared to *stx2c* prophage, however, the *stx2a*-encoding prophage found in 11 of the isolates in this study exhibited a greater diversity both based on Mash distance and whole prophage alignment. The *stx2a*-encoding prophage from each of the lineages causing severe clinical disease in the UK were all distinct, including the two UK sub-lineages (Ia and I/IIa) circulating concurrently and causing outbreaks of HUS in the early 1980s [2,38] (Figure 2). Throughout the 1980s, the number of sub-lineage Ia strains (mainly PT1 and PT4) declined and a new sub-lineage, I/IIb, PT49 emerged. The *stx2a* in the emerging sub-lineage I/IIb PT49 strain was encoded on a bacteriophage that was again distinct from either of the two *stx2a*-encoding prophages found in the representative isolates from the early contemporary sub-lineages Ia and I/IIa. Comparisons between the *stx2a*-encoding prophage in sub-lineage I/IIb revealed similarity to the prophage commonly found to encode *stx2c* (Figures 2 and 4). Furthermore, sub-lineage I/IIb *stx2a* encoding prophage had the same SBI as sub-lineage I/IIa *stx2c* encoding prophages, specifically the *sbcB* gene.

During the 1990s, all three of the dominant 1980s sub-lineages (Ia, I/IIa and I/IIb) declined as a cause of human gastrointestinal disease, and a new sub-lineage emerged. STEC O157:H7 stx2c PT32 belonging to sub-lineage Ic had been circulating in UK and Irish cattle populations for many decades but had not been linked to cases of human disease [5]. However, following acquisition of a *stx2a*-encoding prophage (into the SBI *argW*) which resulted in a change in prophage type to PT21/28 [5,34], sub-lineage Ic became the most common STEC O157:H7 sub-lineage causing gastrointestinal disease and HUS in humans in the UK for the next two decades. The *stx2c*-encoding prophage in lineage Ic had high sequence similarity to *stx2c*-encoding prophage in the other isolates analysed in this study and shared the same SBI, *sbcB* (Table 1, Figures 1 and 4). However, the *stx2a*-encoding prophage acquired by sub-lineage Ic once again differed from those found in the three sub-lineages circulating in the previous decade (Table 1, Figures 1 and 5).

Recently, in the UK there has been a decrease in the number of cases caused by STEC O157:H7 belonging to sub-lineage Ic, and an emergence of sub-lineage IIb PT8 that appears to be associated with the acquisition of a prophage encoding *stx2a* [9]. Strains belonging to this sub-lineage have caused foodborne outbreaks linked to contaminated mixed leaf salad, lamb-based meat products including sausages and mince [39], and an environmental exposure linked to participation in a mud-based obstacle event [39]. Like the *stx2a*-encoding prophage described in sub-linage I/IIb, the *stx2a*-encoding prophage in sub-lineage IIb was similar to the *stx2c*-encoding prophage, and likely the result of horizontal exchange of the *stx2a* gene into a previously *stx2c*-encoding prophage. This is also corroborated by the *stx2a-*encoding prophage in sub-lineage IIb integrating at *sbcB* associated with *stx2c*-encoding prophages (Table 1, Figures 2, 4 and 5).

Importation of STEC O157:H7 strains from outside the UK via contaminated food products are a constant threat. In 2016, a large national outbreak of STEC O157:H7 *stx2a/stx2c* PT34 belonging to sub-lineage IIa occurred in the UK [40], Epidemiological investigations concluded that contaminated red Batavia salad leaves from a non-domestic source was the most plausible vehicle of infection. Analysis of the ONT data from the outbreak strain demonstrated that the *stx2a*-encoding prophage was different from all the *stx2a*-encoding prophage identified in the five major UK sub-lineage. However, this prophage shared sequence similarity with the *stx2a*-encoding prophage in STEC O157:H7 PT32 belonging to sub-lineage Ic, associated with cases of severe gastrointestinal disease in Ireland [5] (Figure 5). Unlike the previously described *stx2a*-encoding prophage, the *stx2a*-encoding prophage in both of these strains share the SBI *yecE*. This prophage also had similarity to the *stx2a*-encoding prophage found in a strain of STEC O55:H7 causing recurrent, seasonal outbreaks of HUS in England [41].

### Summary

Currently, the application of ONT technology for extensive characterisation of STEC O157:H7 genomes at PHE is still under development and therefore, the number of sequences analysed in this study was limited. *stx2a*-encoding prophage exhibited a higher level of diversity and there was little evidence of geographical or temporal patterns of relatedness, or of intra-UK transmission of *stx2a*-encoding prophage between indigenous strains. The *stx2a*-encoding prophage in the UK lineages associated with severe disease therefore appear to be acquired independently and most likely from different geographical and/or environmental sources. These data provide supporting evidence for the existence of a dynamic environmental reservoir of *stx2a*-encoding prophage that pose a threat public health due to their potential for integration into competent, indigenous sub-lineages of *E. coli* O157:H7. Finally, we provide further evidence that, compared to *stx2a*-encoding prophage, *stx2c*-encoding prophage exhibit a high level of similarity across lineage, geographical region and time, and have likely been maintained and inherited vertically.

## 8. Author statements

### 8.1 Authors and contributors

TJD and CJ conceptualised the project. DRG performed DNA extractions, library preparations and sequencing of isolates. DAY performed data processing, genome assembly, genome polishing, genome annotation. DRG and DAY and created the Easyfig diagrams. DRG performed prophage comparison using Mash and TJD wrote associated scripts. DAY, DRG, TJD and CJ wrote the original manuscript. DAY, DRG, TJD, CJ, DLG and SEG reviewed and edited the manuscript. TJD, CJ and DLG supervised DRG. DRG, TJD and CJ supervised DAY.

